# The role of Selenoprotein N in the differentiation of erythroid progenitors during stress erythropoiesis

**DOI:** 10.1101/506345

**Authors:** Robert F. Paulson, Shaneice Nettleford, Chang Liao, Yuanting Chen, Siyang Hao, Mireille Baltzinger, Mélanie Thami-Braye, Alain Lescure, K. Sandeep Prabhu

**Affiliations:** Department of Veterinary and Biomedical Sciences, Penn State Cancer Institute, The Pennsylvania State University, 115 Henning Building, University Park, PA 16802, USA; Université de Strasbourg, CNRS, Architecture et Réactivité de l’ARN, UPR 9002, IBMC-15 rue René Descartes, F-67000, Strasbourg, France

## Abstract

Low serum Se is independently associated with anemia in elderly population, dialysis patients, sickle-cells patients, and hypothyroidism patients. Previous work from our laboratory showed that dietary Se deficiency in mice showed mild anemia indicating activation of stress-erythropoietic mechanisms. Unlike steady state erythropoiesis that is primarily responsible for homeostasis to produce new erythrocytes at a constant rate, stress erythropoiesis predominates when the bone marrow cannot generate sufficient erythrocytes. During such a process, short-term reconstituting hematopoietic stem cells (CD34^+^Kit^+^Sca1^+^Lin^neg^) migrate to the spleen leading to the proliferation and differentiation of stress-erythroid progenitors (SEPs). These cells lead to stress burst forming unit-erythroid cells (BFU-E) followed by terminal differentiation to erythrocytes. Recent studies demonstrate deficits in selenoproteins block the expansion and development of stress BFU-E with defects in terminal differentiation. Analysis of selenoprotein expression showed that selenoprotein W (SelenoW) was highly expressed in developing SEPs. CRISPR-Cas9 knockout of SelenoW blocked the proliferation of immature SEPs in murine and human stress erythropoiesis cultures demonstrating a central role for SelenoW in stress erythropoiesis. Using the two-culture system to generate SEPs, selenoprotein N (SelenoN) expression increased as the progenitors transition from self-renewing “stem cell like” progenitors to form committed erythroid progenitors. SelenoN^−/−^ mice showed significantly slower erythroid recovery following phenylhydrazine (PHZ)-induced acute hemolytic anemia. As in the muscle satellite cells where SelenoN regulates cellular Ca^2+^ signaling, SelenoN may also regulate Ca^2+^ signaling in SEPs to modulate pathways of differentiation. In summary, these data suggest that multiple selenoproteins, including SelenoN and SelenoW, coordinately regulate stress erythropoiesis.

## Introduction

Anemia is a significant human health problem caused by multiple etiologies. Intrinsic defects in erythropoietic progenitors can impair hemoglobin production and the generation of new erythrocytes as observed in sickle cell anemia and thalassemia. In addition, many anemias present as a secondary pathology as observed in the anemia of inflammation and anemias caused by nutritional deficiencies (1–5). Although one of the leading causes of anemia in the United States and worldwide is the nutritional deficiency of iron, deficiencies in other trace elements are associated with anemia have been characterized (6). Low serum Se was also independently associated with anemia among older men and women in the USA and elsewhere (7–9). Dialysis patients showed significantly lower plasma Se levels that correlated with decreased half-lives of erythrocytes and thrombocytes compared to healthy controls (10). In sickle cell anemia patients, Se deficiency was associated with decreased expression of the highly Se-responsive selenoprotein glutathione peroxidase-1 (GPX1) in addition to a weakened antioxidant potential (11). A causal association between hypothyroidism and anemia (12) and subclinical hypothyroidism, due to Se deficiency (and decreased selenoprotein deiodinases), was associated with lower hemoglobin levels (13). Altogether, these studies suggest a key role for Se in erythropoiesis during anemia. The standard treatments for anemia include whole blood transfusion and administration of recombinant human erythropoietin (rhEpo). However, transfusion therapy poses problems in the treatment of chronic anemia with increased risk of allo-immunization and the potential to induce inflammation or infection. rhEpo is not effective in all types of anemia and can compromise chemotherapy(5, 14–23). Therefore, the development of new treatments for anemia is a significant unmet need in the field.

One approach to identifying new therapies is to better understand the physiological response to anemic stress. Steady state erythropoiesis is primarily homeostatic and produces new erythrocytes at a constant rate. However, when the bone marrow cannot generate sufficient erythrocytes, stress erythropoiesis predominates (24). Stress erythropoiesis is best understood in mice where it occurs extra-medullary in the adult spleen and liver (24–28). Unlike steady state erythropoiesis, stress erythropoiesis utilizes a set of progenitors and signals that are distinct from the ones used in bone marrow erythropoiesis (29, 30). Steady state erythropoiesis produces erythrocytes at a constant rate; however, the strategy of stress erythropoiesis is different. Stress erythropoiesis proceeds through four stages. In the initial stage, short-term reconstituting hematopoietic stem cells (CD34^+^Kit^+^Sca1^+^Lin^neg^) migrate to the spleen where Hedgehog and Bmp4 expressed in the spleen microenvironment specify the cells as stress-erythroid progenitors (SEPs), a set of erythroid restricted cells (29). During the second stage, anemic stress induces the rapid proliferation of these cells in the spleen, but they maintain their stem cell markers and retain their undifferentiated state (27, 31, 32). An influx of erythropoietin (Epo) marks the third stage which induces a transition from proliferating stem cell like progenitor to committed erythroid progenitor. At this point cells acquire the ability to differentiate(31, 32). During the final stage, the stress erythroid progenitors synchronously differentiate to generate a wave of new erythrocytes that maintain erythroid homeostasis until the bone marrow can resume production. It is expected that a better understanding of the mechanisms that regulate stress erythropoiesis would identify new targets for therapeutic intervention to treat anemia without relying on transfusion or injection of rhEpo.

Selenoproteins function as redox gatekeepers in cells via the active site Sec residue. In addition to the hierarchical expression pattern, selenoproteins possess varying degrees of peroxidase activity(33). Some selenoproteins also contain a thioredoxin-like fold with a conserved CxxU (C: Cys; U: Sec) motif that is critical for its function as a disulfide reductase in concert with other redox couples, which is preceded by an interaction with cellular protein substrates. Erythropoiesis and erythrocytes are particularly prone to oxidative stress due to the presence of iron, heme and O_2_. Mutation of *Trsp*, which encodes the tRNA^[Sec]^, abolishes selenoprotein production and is embryonic lethal(34). However, conditional deletion in the hematopoietic system leads to increased oxidative stress and induction of the Nrf2-dependent antioxidant pathway. Double mutants in Trsp and Nrf2 exhibit an exacerbated anemia characterized by increased levels of H_2_O_2_ in erythroblasts demonstrating that increased oxidative stress profoundly affects erythroid development(35). Previous work from our laboratory showed that dietary Se deficiency leads to increased hemoglobin oxidation and formation of Heinz bodies(36). Se-deficient mice exhibit a mild anemia. Recently we demonstrated that Se deficiency or lack of selenoproteins due to conditional deletion of *Trsp* in hematopoietic cells severely compromised stress erythropoiesis at several levels(37). Deficits in selenoproteins blocked the expansion and development of stress burst forming unit-erythroid cells (BFU-E) and caused defects in terminal differentiation. Analysis of selenoprotein expression showed that selenoprotein W (SelenoW) was highly expressed in developing SEPs. CRISPR-Cas9 knockout of SelenoW blocked the proliferation of immature stress progenitors in murine and human stress erythropoiesis cultures demonstrating a central role for SelenoW in stress erythropoiesis(37).

Selenoprotein N (SelenoN) is another Sec containing protein localized in the endoplasmic reticulum (ER) lumen, ubiquitously expressed in all tissues. However, loss-of-function of the SELENON gene leads to a group of neuromuscular disorders referred to as SelenoN(SEPN1)-related myopathies characterized by trunk muscle atrophy, scoliosis, and respiratory defects (38). This muscle specific phenotype in human and studies on *SelenoN*^−/−^ mouse suggested that, despite its widespread expression, this selenoprotein is particularly important in molecular processes relevant to muscle development and maintenance(38). Interestingly, decreased SelenoW expression was also linked to the development of a muscle phenotype in cattle, the white muscle disease. SelenoN^−/−^ mouse suggested that this selenoprotein essential for muscle regeneration and satellite cell maintenance in mice and humans (39). While the function of SelenoN has been elusive, recent breakthroughs have indicated that SelenoN protects the ER from hydroperoxides generated by the endoplasmic reticulum oxidoreductin-1 (ERO1), an ER thiol oxidase, to restore calcium homeostasis (40). SelenoN was reported to interact with the sarco/endoplasmic reticulum Ca^2+^-ATPase (SERCA) pump via its thioredoxin motif indicating a redox-dependent enhancement of its function in Ca^2+^ homoestasis (40). Ca^2+^ signaling regulates the response to ER stress through the activation of Perk kinase (Eif2aK3), which plays a key role in regulating muscle satellite cell quiescence and self-renewal(41, 42). Interestingly, SelenoN^−/−^ mice exhibited a defect in muscle satellite cells and *SelenoN*^−/−^ mice have fewer satellite cells and progressively lose satellite cells over repeated muscle injuries, which impairs their ability to maintain regenerative cycles (39). Stress erythropoiesis also utilizes a self-renewing population progenitor cells. Similarly, a role for Perk signaling is supported by the observation that ATF4^−/−^ mice have defect in stress erythropoiesis(43). ATF4 translation is stimulated by Ca^2+^ dependent Perk signaling(44, 45). These observations suggest that similar to muscle satellite cells, SelenoN may also regulate Ca^2+^ signaling in SEPs. Here we show that SelenoN expression increases as the progenitors transition from self-renewing “stem cell like” progenitors to committed erythroid progenitors and examine the role of SelenoN in the erythroid recovery from phenylhydrazine (PHZ)-induced acute hemolytic anemia.

## Materials and Methods

C57BL/6 mice of both sexes were maintained on unrestricted access to lab chow and water. Approximately eight to 12-week old mice were used to isolate bone marrow. For the PHZ challenge experiment, eight to ten-months old *SelenoN*^−/−^ male mice on a C57BL/6 background were used and compared to *SelenoN*^+/−^ littermates. All procedures were performed under guidelines from the Animal Use and Care Committee at Pennsylvania State University.

### Phenylhydrazine treatment

Mice were injected intraperitoneally with PHZ at a dose of 100 mg/kg body weight as previously described (37). Peripheral blood was collected for hematocrit measurement, using heparinized 75μL microhaematocrit capillary tubes. Capillary tubes were centrifuged at 500g for 10 min and both total blood volume and blood cell pellet were measured. Percentage hematocrit was defined as the ratio of blood cells over total blood volume.

### Ex-vivo *culture of SEPs*

Bone marrow from WT C57BL/6 mice were cultured in the two-phase culture system as described recently (37). Briefly, bone marrow cells were cultured in the stress erythroid expansion media (SEEM), which comprised of Iscove’s Modified Dulbecco’s Media (IMDM) containing 1% (m/v) BSA, 0.0007% (v/v) β-mercaptoethanol, 10% (v/v) FBS, 4mM L-glutamine, Insulin (10μg/mL), Transferrin (200μg/mL), Scf (50ng/mL), Bmp4 (15ng/mL), Gdf15 (30ng/mL), and Shh (25ng/mL), for seven days to enrich for SEPs. Following expansion, the above cells were shifted to stress erythroid differentiation media (SEDM), which is SEEM supplemented with recombinant erythropoietin (Epo; 3U/mL), and the cultures grown in hypoxia conditions (1% O_2_, 5% CO_2_, 37°C) for three days. Cells were isolated from the SEDM media on day1 and day 3, referred to as D1 and D3, respectively, and used in qPCR assays.

### Quantitative polymerase chain reaction (qPCR) and gene expression analysis

RNA isolation from cells or tissues was performed using TRIzol reagent (Life Technologies) according to the manufacturer’s instructions. cDNA was synthesized by using High Capacity cDNA Reverse Transcriptase kit (Applied Biosystems). qPCR reactions were performed with SYBR™ green master mix (Quanta Biosciences). Primer sequences are as follows:

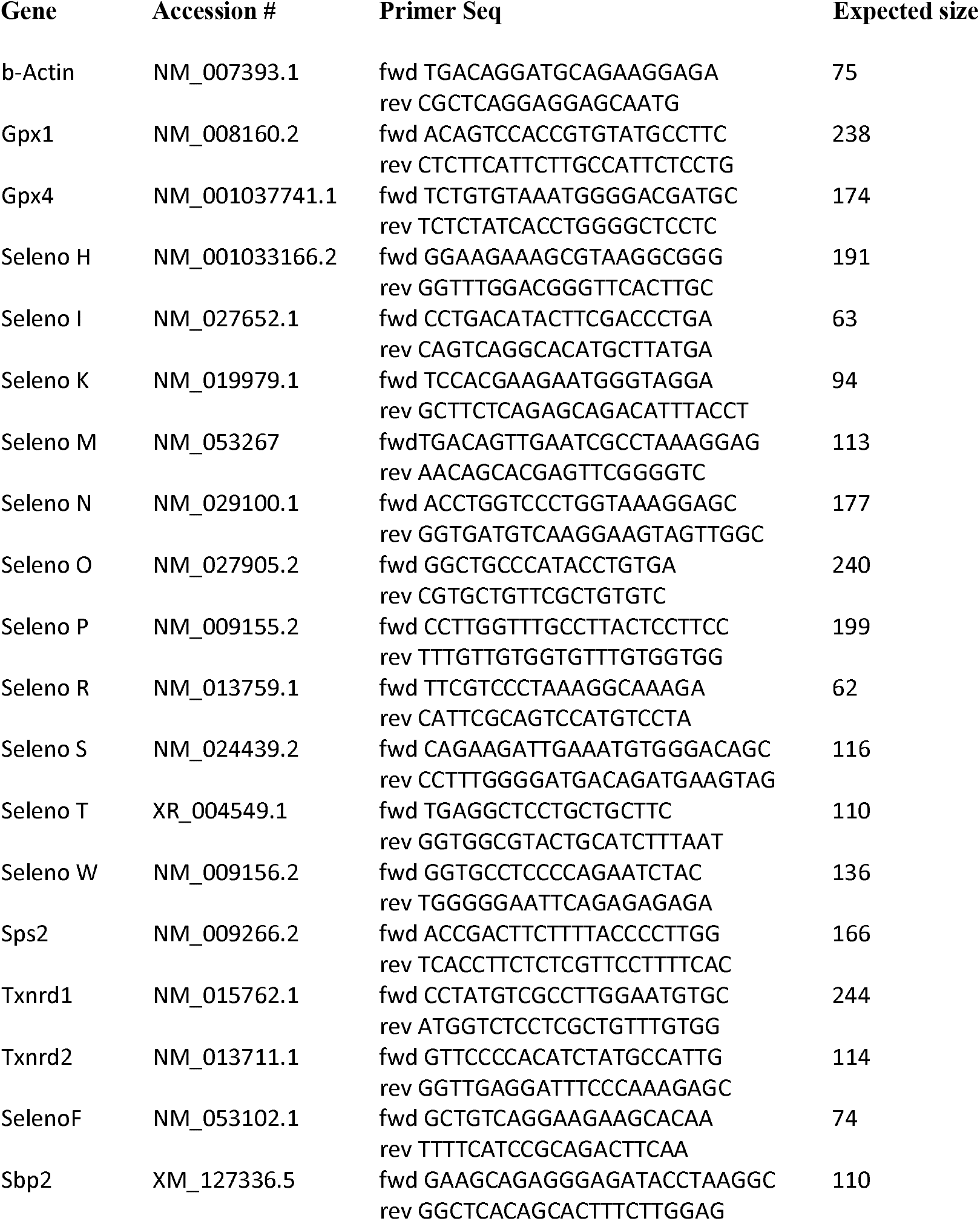

## Results and Discussion

SEPs develop from short-term reconstituting hematopoietic stem cells (CD34^+^Kit^+^Sca1^+^Lin^−^; ST-HSCs) that migrate to the spleen and adopt a stress erythroid fate (28, 29). The proliferation and transition to differentiation must be coordinated to ensure that sufficient new erythrocytes are produced. Our recently published data show that dietary Se deficiency or mutations that block the production of selenoproteins severely compromise stress erythropoiesis (37). We observed defects at multiple levels. Immature SEPs failed to proliferate in-vivo and in-vitro along with defective development and differentiation of BFU-Es.

The transition from proliferating SEPs to committed erythroid progenitors capable of forming BFU-E colonies is regulated by Epo signaling. We have developed an in vitro culture system that recapitulates the events in vivo and generates functional SEPs that are transplantable(32). Mutation of ATF4 leads to defects in the development of stress BFU-E, which suggests that ER stress may play a role in regulating stress erythropoiesis(43). Translation of ATF4 is induced by Ca^2+^ dependent activation of Perk kinase(44, 45). We examined the change in expression of ER-resident selenoproteins that play a key role in Ca^2+^ homeostasis in SEPs on days 1 and 3 after Epo was added to the media to induce the transition to differentiation. Analysis of selenoprotein expression in SEPs showed that SelenoN was significantly upregulated when cultures were switched from expansion media (SEEM) to Epo containing differentiation media (SEDM). In addition to SelenoN, only Txnrd2 and SelenoO showed statistically significant changes in expression at the transcript level (Fig 1).

**Figure 1:**
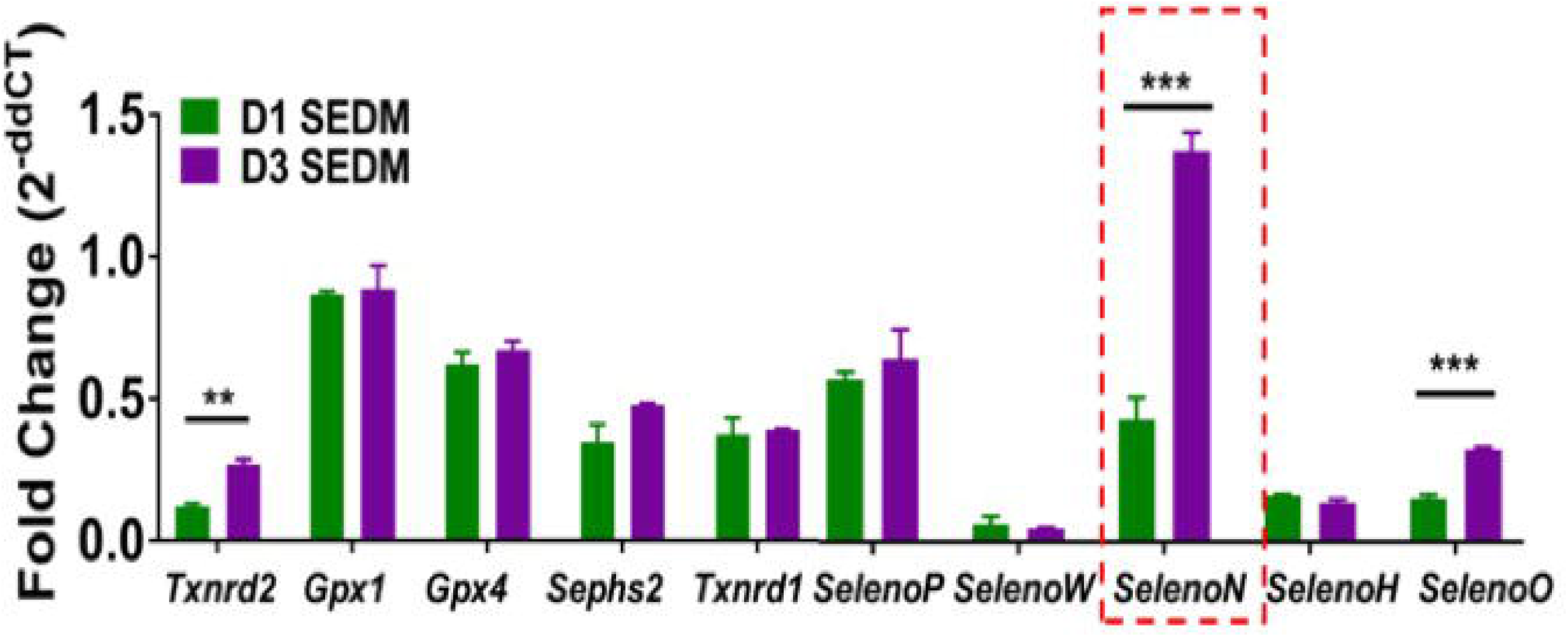
Expression of selenoproteins during SEP differentiation cultures. Mouse bone marrow was cultured in SEEM for seven days followed by SEDM for one day (D1) or three days (D3). Expression of selenoproteins was determined by quantitative PCR. All samples were normalized to the expression of Gapdh. The data was compared to cells cultured for seven days in SEEM media. Data shown is n = 3 per time point; *, **, ***: p<0.05; p<0.01; p<0.005, respectively.

Txnrd2 is a pyridine nucleotide-disulfide oxidoreductase and a part of the thioredoxin (Trx): thioredoxin reductase redox couple. Unlike Txnrd1 that is cytoplasmic, Txnrd2 is a mitochondrial enzyme that scavenges reactive oxygen species during ER stress and is also known to suppress the therapeutic efficacy of proteasome inhibitors such as bortezomib (Velcade) (46). In multiple myeloma, bortezomib activates Kruppel-like Factor 9 (KLF9) that transcriptionally represses Txnrd2 (47). However, in non-cancer cells, the control of Txnrd activity could have important ramifications. When ER homeostasis is disturbed, mitochondrial dysfunction could be disrupted thereby making Txnrd2 and other antioxidant proteins crucial in cell fate decisions, particularly in erythrocyte development. Another selenoprotein that was differentially regulated was SelenoO. Although not much is known about its biological function, SelenoO is also a mitochondrial protein that has been recently reported to function as a pseudokinase, which catalyzes the AMPylation of proteins involved in redox homeostasis (48). Interestingly, SelenoO deficiency was reported to affect chondrocyte differentiation and suggested its likely role in endemic osteoarthropathy that is commonly seen during Se deficiency(49). However, there is no literature regarding the role of SelenoO in erythrocyte development.

Given that SelenoN is highly regulated during differentiation and considering its role in the redox-dependent calcium transport control by the SERCA pump, we further tested the ability of *SelenoN*^−/−^ mice to respond to stress erythropoiesis during PHZ administration. Our experiments showed that *SelenoN*^−/−^ mice treated with PHZ (100 mg/kg) led to a significant decrease in hematocrit levels compared to the heterozygous (*SelenoN*^+/−^) mice. Furthermore, *SelenoN*^−/−^ mice exhibited a slower recovery during the 10-day period post PHZ treatment (Fig. 2). As expected *SelenoN*^−/−^ did not show as severe phenotype as the complete loss of selenoproteins as observed in the *Trsp*^−/−^ mice. These data suggest that multiple selenoproteins, including SelenoN and SelenoW, coordinately regulate stress erythropoiesis. These data provide further justification for detailed molecular analyses to delineate the role of SelenoN and other selenoproteins in the regulation of SEP differentiation and erythrocyte development.

**Figure 2:**
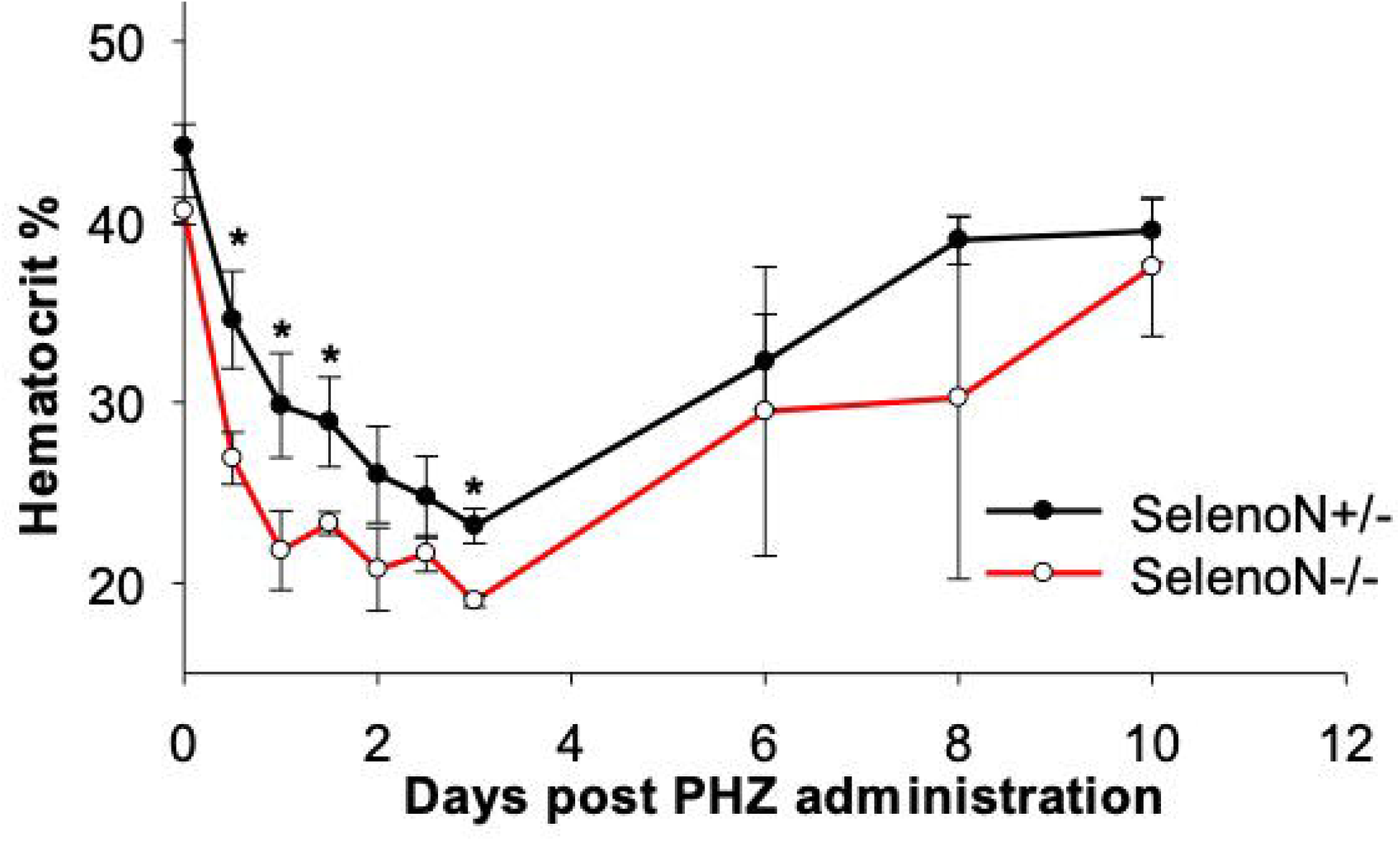
Analysis of hematocrit during recovery in *SelenoN*^+/−^ and *SelenoN*^−/−^ mice upon treatment with 100% PHZ. Age and sex-matched SelenoN^+/−^ and SelenoN^−/−^ mice were administered phenylhydrazine (PHZ) at 100 mg/kg body weight and the hematocrit were followed during 10 days post PHZ administration. Data shown (mean hematocrit ± SEM) is n = 4-5 per genotype; *, p<0.05

